# How sex-biased dispersal affects conflict over parental investment

**DOI:** 10.1101/026732

**Authors:** Bram Kuijper, Rufus A. Johnstone

**Affiliations:** CoMPLEX, Center of Mathematics and Physics in the Life Sciences and Experimental Biology, University College London, Gower Street, WC1 6BT, London, United Kingdom; Department of Genetics, Evolution and Environment, University College London, Gower Street, WC1 6BT, London, United Kingdom; Behaviour and Evolution Group, Department of Zoology, University of Cambridge, Downing Street, CB2 3EJ Cambridge, United Kingdom

## Abstract

Existing models of parental investment have mainly focused on interactions at the level of the family, and have paid much less attention to the impact of population-level processes. Here we extend classical models of parental care to assess the impact of population structure and limited dispersal. We find that sex-differences in dispersal substantially affect the amount of care provided by each parent, with the more philopatric sex providing the majority of the care to young. This effect is most pronounced in highly viscous populations: in such cases, when classical models would predict stable biparental care, inclusion of a modest sex difference in dispersal leads to uniparental care by the philopatric sex. In addition, mating skew also affects sex-differences in parental investment, with the more numerous sex providing most of the care. However, the effect of mating skew only holds when parents care for their own offspring. When individuals breed communally, we recover the previous finding that the more philopatric sex provides most of the care, even when it is the rare sex. Finally, we show that sex-differences in dispersal can mask the existence of sex-specific costs of care, because the philopatric sex may provide most of the care even in the face of far higher mortality costs relative to the dispersing sex. We conclude that sex-biased dispersal is likely to be an important, yet currently overlooked driver of sex-differences in parental care.

## 1 Introduction

Although mothers and fathers share a common genetic stake in the survival of their young, their evolutionary interests are rarely fully aligned [1–3]. Each parent typically stands to gain if the other bears more of the costs of raising the young, leading to sexual conflict over the provision of parental care [4–7]. Such conflict is thought to be an important driver of between-species variation in forms of care [8] and is associated with a range of conspicuous behaviours, such as brood concealment [9], infanticide [10, 11], brood desertion [12, 13] and negotiation between parents over care [e.g., 14–16].

Over the years, a substantial body of theory has been developed to explain the evolution sex-differences in parental care [for reviews see 6, 7], focusing chiefly on behavioural interactions within the family (e.g., negotiation or coercion between parents [17–23]) or between parents and helpers [24, 25]). This emphasis on interactions at the family level can perhaps be attributed to the self-contained nature of the family-unit, which allows researchers to concentrate on the behaviour of a small number of family members in the context of a clearly defined “nursery” environment [26]. However, the downside of such a narrow focus is that the potential impact of processes at the population level has been largely ignored in existing models of parental investment. So far, the only population-level process that has received much attention is the feedback between parental investment and mate availability [8, 27–29]. The possible influence of other population-scale processes on the provision of care remains poorly understood.

Dispersal is one such population-level process, which is thought to influence the resolution of other forms of family conflict (e.g., genomic imprinting: [30–32], parent-offspring conflict [33] and infanticide [34]). The importance of dispersal in the familial context raises the question whether it might also affect the outcome of sexual conflict between parents over care. A key hint that dispersal may influence investment decisions comes from models that focus on more abstract social behaviour (e.g., helping and harming) [35, 36]. These models show that sex-differences in dispersal can favour substantial asymmetries in helping behaviours between the two sexes. Given that sex-biases in dispersal are common in the animal kingdom [37–39], it seems likely that patterns of dispersal might provide useful insights into the evolution of uniparental and biparental care.

Here, we set out to examine the impact of sex-biased dispersal on parental care, in particular on sex-differences in parental investment. We build on the classical theoretical model of biparental care by Houston & Davies [17] embedding it in a demographically explicit framework familiar from recent models of social evolution in structured populations [35, 40–43].

## 2 The model

### A demographical model of parental care

Our approach is to develop an evolutionary demographic model [40, 44–47] of the provision of care to a brood by male and female parents.

Consider a population distributed over infinitely many territories [48, 49], in which individuals reproduce sexually and generations are overlapping. Each territory contains *n*_f_ adult female breeders and *n*_m_ adult male breeders. Time proceeds in a series of discrete breeding seasons, during each of which a given female mates at random within her patch to produce a large number of offspring, in a 1: 1 sex ratio. We assume that all males on the patch have identical mating success. When *n*_f_ = *n*_m_, this could entail females and males on a patch pairing up at random, with each female’s offspring all being fathered by the same male, or each of a female’s offspring might be fathered by a randomly chosen male; both scenarios give identical results. When *n*_f_ ≠ *n*_m_, the latter interpretation is more natural.

The individual parental efforts of males and females are given by *u*_m_ and *u*_f_ respectively, and offspring survival, which depends upon the combined efforts of both parents, is given by *b*(*u*_f_ + *u*_m_). Following classical models of parental investment [4, 5, 19, 50, 51], we assume that *b* increases, at a diminishing rate, with an increase in either parent’s level of effort *u*_*x*_, *x* ∊{m, f} (so that *∂b/∂u*_*x*_ > 0 and 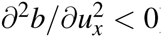 In addition, an increased level of parental effort *u*_*x*_ is assumed to come at a cost to the parent’s future fitness, through an increase in its mortality probability *µ*_*x*_ Ξ µ_*x*_ (*u*_*x*_) [4]. Again, in accordance with previous models (e.g., [19, 21]), we assume that parental mortality increases in accelerating fashion with an individual’s effort *∂µ*_*x*_*/∂u*_*x*_ > 0 and 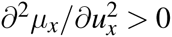.

Of all offspring produced during each time step that survive, a fraction *h*_m_ of sons and a fraction *h*_f_ of daughters remain on the natal patch, while fractions *d*_m_ = 1 − *h*_m_ and *d*_f_ = 1 − *h*_f_ disperse to other randomly chosen patches. After dispersal, offspring on a patch (both native and immigrant) compete for any breeding vacancies created by the mortality of adult members of their own sex. The offspring that fail to claim a breeding vacancy die, after which the same sequence of events is repeated in the next time step.

### Selection gradients

To model the evolution of male and female parental care, we use an adaptive dynamics approach [40, 52, 53]. This method assumes that evolutionary change in female and male care levels *u*_f_ and *u*_m_ occurs through the successive invasion and substitution of mutations of slight effect. In the Appendix, we derive fitness expressions *W* (*u*_f_ + *δu*_f_; *u*_f_, *u*_m_) (or *W* (*u*_m_ + *δu*_m_; *u*_f_, *u*_m_)) for the fitness of a mutant female (or male) with care level *u*_f_ + *δu*_f_ (*u*_m_ + *δu*_m_) in a resident population that has female care levels *u*_f_ and male care levels *u*_m_. When mutations in male and female care levels occur independently (i.e., no pleiotropy), the rate and direction of evolutionary change in *u*_f_ and *u*_m_, is then proportional to the selection gradients *∂W* (*u*_f_ + *δu*_f_; *u*_f_, *u*_m_) */∂δu*_f_ and *∂W* (*u*_m_ + *δu*_m_; *u*_f_, *u*_m_) */∂δu*_m_ evaluated at *δu*_f_ = *δu*_m_ = 0. Expressions for these selection gradients are derived using a neighbour-modulated fitness (also called direct fitness) approach [45, 46, 54] in the Appendix. Consequently, we can solve for the equilibrium levels of female and male care (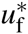 and 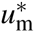) at which both selection gradients vanish.

### Individual-based simulations

We also ran individual-based simulations to corroborate our analytical results. We simulated a population of 2000 patches, each containing *n*_f_ female breeders and *n*_m_ male breeders. Each individual bears two unlinked, diploid loci *u*_f_ and *u*_m_. Alleles at each locus interact additively. Each allele mutates with a per-generation probability of *ν* = 0.01. In case a mutation occurs, a random number is drawn from a normal distribution with mean zero and variance 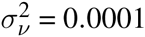and added to current allelic value. During each time step, a female chooses a random partner and produces *M·b*(*u*_f_ + *u*_m_) offspring, where *M* = 200. Male and female offspring disperse to a random patch with probability *d*_m_ or *d*_f_ or remain at the natal patch with probability 1 − *d*_m_ and 1 − *d*_f_. After the offspring production stage, female and male breeders die with respective probabilities *µ*_m_(*u*_m_) and *µ*_f_(*u*_f_). In case of a death, a new offspring is randomly sampled from the pool of immigrant and local offspring in each patch, after which the next time step begins. Simulations ran for 100 000 generations. The simulations are coded in C and are available from the first author’s website.

### Male and female care in well-mixed populations

We first focus on a simple ‘baseline’ scenario in which the number of breeding adults per patch is identical for both sexes, *n* = *n*_f_ = *n*_m_, and in which males and females care only for their own offspring (i.e., no communal breeding [55, 56]), so that the survival of an individual offspring depends only upon the efforts of its own father and mother. For a well-mixed population (*h*_f_ = *h*_m_ = 0), we then find that the equilibrium parental efforts (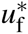 and 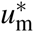) of mothers and fathers satisfy:

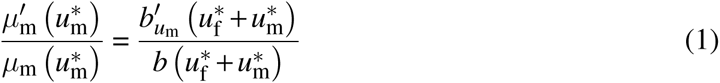

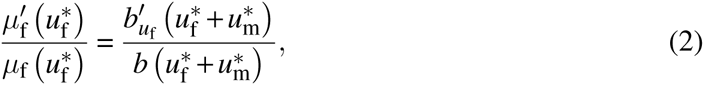

where 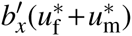 reflects the partial derivative of juvenile reproductive value with respect to the variable *x*. In other words, for each parental sex, the proportional increase in parental mortality due to a slight increase in effort (corresponding to the left-hand side of the equations) must precisely balance the proportional increase in offspring survival (corresponding to the right-hand side of the equations).

### Sex-specific care when dispersal is limited

Again, we focus on a scenario where parents care only for their own offspring (no communal breeding). We now consider a viscous population in which dispersal is limited (*h*_f_, *h*_m_ > 0) and where the number of male breeders *n*_m_ is not necessarily identical to the number of female breeders *n*_f_ (mating skew). Dependent on the level of mating skew, males sire, on average, offspring from multiple females and hence provide care for multiple broods (when males are rare, *n*_m_ *< n*_f_) or sire only part of a brood of a single female (when males are common, *n*_m_ > *n*_f_) and thus only care for that part of the brood. As the offspring survival function *b*(*u*_f_ + *u*_m_) measures effort at the level of a single brood, we thus need to weigh a male’s contribution by his expected number of mates *n*_f_*/n*_m_, so that we have 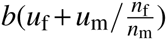. Incorporating these assumptions in a demographical model of parental care set out in the Supplement, we show that equilibrium female and male parental effort (eqns. [S24, S25]) now satisfy:

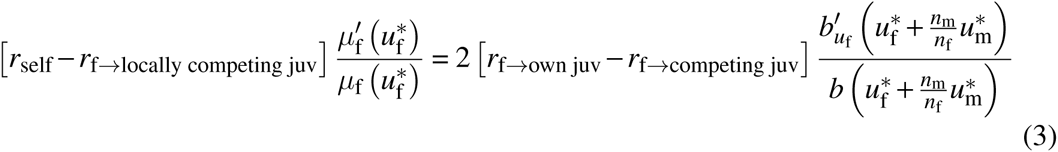

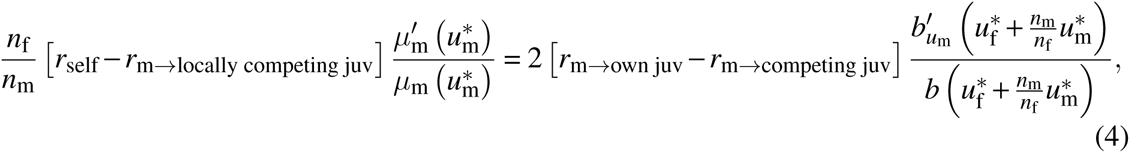

Here, terms on the left-hand side show that the proportional increase in parental mortality due to a slight increase in effort by each sex is weighted by the difference between *r*_self_, which denotes the relatedness of the focal breeder to itself (*r*_self_ = 1), and *r*_*x*→locally competing juv_, which denotes the relatedness of a focal breeder of sex *x* to locally competing juveniles of the same sex, one of which will replace the focal breeder when it dies. On the right-hand side, the proportional increase in offspring survival due to a slight increase in parental effort is weighted by twice the difference between *r*_*x*→own juv_, which denotes the relatedness of a focal to its own juvenile offspring and *r*_*x*→competing juv_, which denotes its relatedness to offspring competing with its own young (the factor of two appears because in a sexual population every parent produces on average two offspring, one of each sex). Note that these relatedness coefficients depend, in turn, on a number of demographical parameters, including the sex-specific dispersal probabilities *h*_f_ and *h*_m_, the sex-specific mortality rates *µ*_m_(*u*_m_) and *µ*_f_(*u*_f_) and the number of male and female breeders *n*_m_ and *n*_f_. Explicit expressions for these relatedness coefficients are derived in the Supplementary Information, section S1.4.

Note that in a well-mixed population, in which there is no philopatry *h*_f_ = *h*_m_ = 0, we have 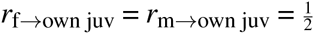, *r*_f→locally competing juv_ = *r*_m→locally competing juv_ = *r*_f_→competing juv =*r*_m→competing juv_ = 0, so that equations (3,4) reduce to eqns. (1,2).

Eqns. (3,4) can be used to investigate the influence of sex-biased philopatry on caring effort, for example by considering a scenario in which males are more philopatric than are females (*h*_m_ > *h*_f_). As a result of such a dispersal asymmetry, a male’s average relatedness *r*_m→local juv_ to local juveniles of his own sex increases compared to a female’s average relatedness *r*_f→local juv_ to juveniles of her own sex. The left-hand side in eq. (4) therefore becomes smaller relative to the left-hand side in eq. (3). One can interpret this as a decrease in the effective mortality cost of care for males compared to females, due to the fact that in dying, a male breeder is more likely than is a female to free up a breeding spot for a related offspring. However, inspecting the right-hand side of eq. (4), *h*_m_ > *h*_f_ also entails a decrease in the marginal benefits of increased male effort. This is because the term *r*_m→competing juv_ becomes larger, as a philopatric male who raises more surviving young is also more likely than is a female to displace related offspring. The balance between these two effects, however, favours greater parental effort by the more philopatric sex (here males), because it can be shown that *r*_m→competing juv_ on the right-hand side is smaller to an order 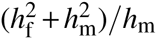 relative to the coefficient *r*_m→local juv_ on the left-hand side (see eqns. S27, S29). As a result, male-biased philopatry leads males to increase their level of parental effort, while females reduce theirs.

### 2.1 Result 1: the philopatric sex provides more care to the brood

Figure 1 provides a simple illustration of the dramatic effect that sex-differences in philopatry can have on the levels of parental effort provided by males and females: in contrast to classical models [17, 19], which predict that both parents should invest equally in their young (Figure 1A), even modest sex-biases in philopatry lead to the more philopatric sex (males in this example) providing the majority of care to young (Figure 1B). Stronger sex-biases in philopatry can even lead to scenarios in which the philopatric sex effectively become sole carers, while the contribution by the dispersing sex to the brood is negligible (Figure 1C). This pattern is robust to variation in absolute sex-specific dispersal probabilities, provided that the direction of sex-specific bias is unchanged, as shown in Figure 2. The effect is most pronounced when a single breeding pair occupies a patch (Figure 2A). With a larger number of breeding pairs per patch, the overall asymmetry may be less pronounced (as relatedness to young is reduced due to the presence of other breeding pairs), but the most philopatric sex still provides the majority of care (Figure 2B).

**Figure 1:**
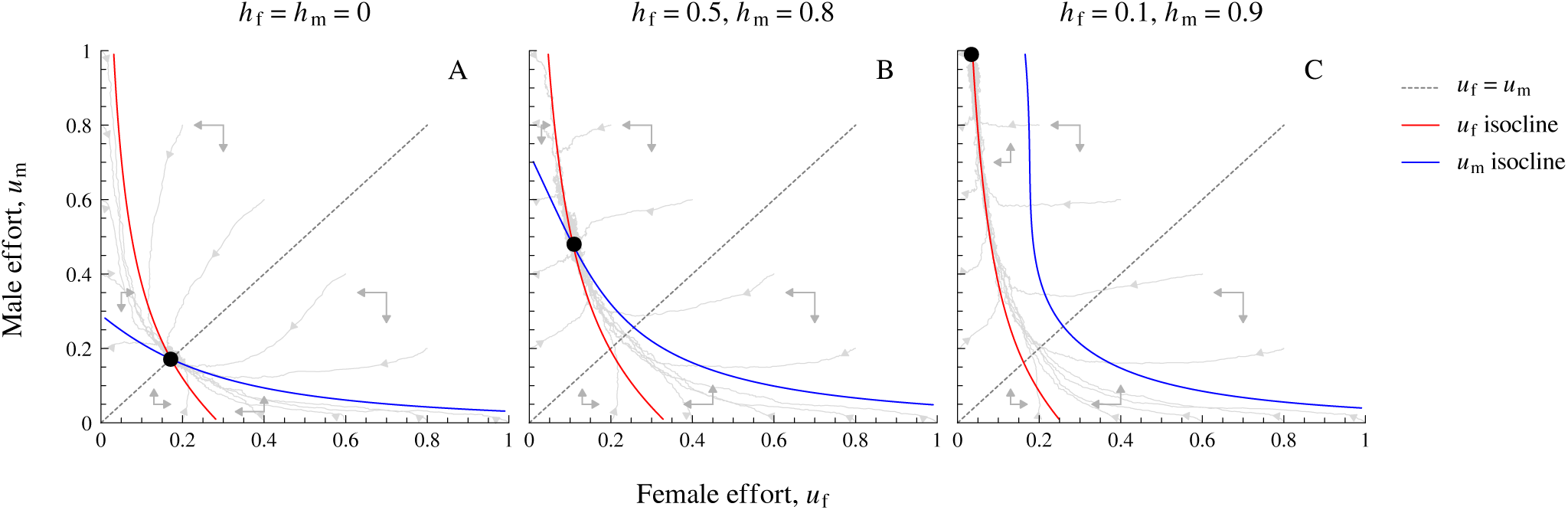
Sex-biases in dispersal can drive evolutionary transitions from biparental to near-uniparental care. The black dot in each panel shows the evolutionary stable level of female 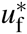 and male parental care 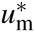for different combinations of male and female philopatry, *h*_m_ and *h*_f_ respectively. Panel A: the baseline case of a well-mixed population with complete migration, in which there are no sex-differences in care, *u*_f_ = *u*_m_. Panel B: when males are more philopatric than females (*h*_m_ > *h*_f_), males provide the majority of care to young. Panel C: when males are extremely philopatric in comparison to females, selection favours males to provide almost all care to young, whereas the amount of female care is negligible. Parameters: *n*_m_ = *n*_f_ = 1, *k*_f_ = *k*_m_ = 0.1. Red line: the selection differential of female effort vanishes, d*W*/d*u*_f_ = 0. Blue line: the selection differential of male care vanishes, d*W*/d*u*_m_ = 0. Light grey lines: individual-based simulations.

**Figure 2:**
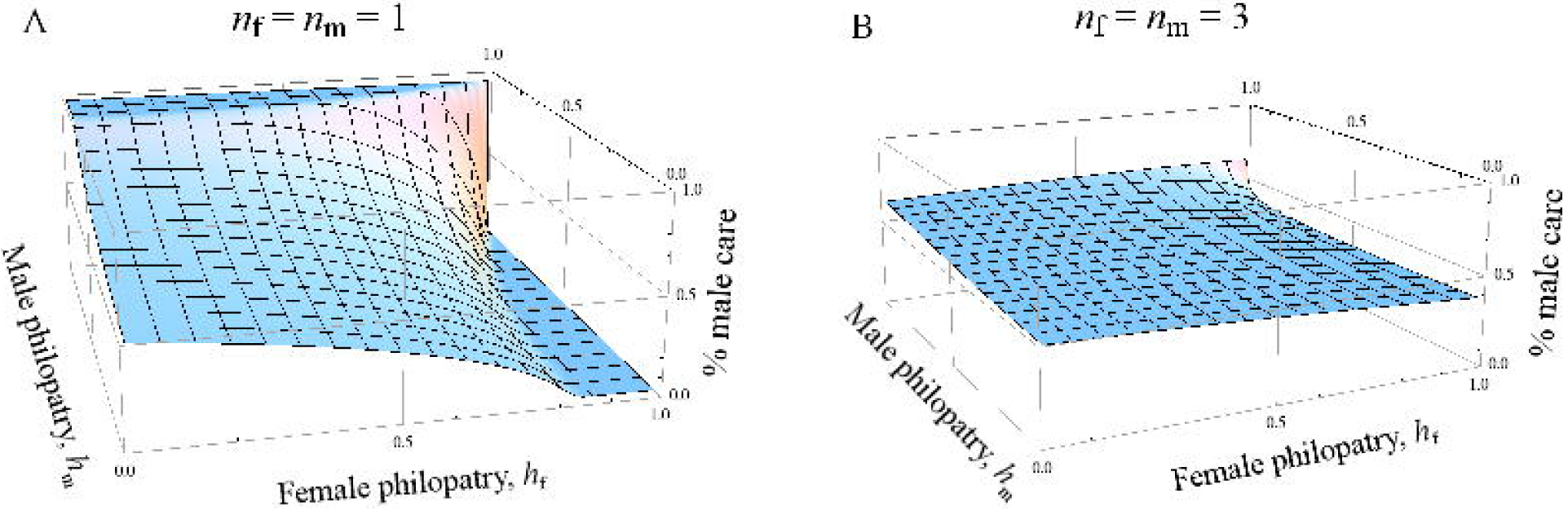
The equilibrium proportion of male care 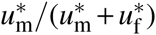as a function of the amount of philopatry in both sexes. Panel A: in highly viscous populations with few individuals per patch (*n*_f_ = *n*_m_ = 1), slight sex-biases in philopatry cause pronounced sex-differences in care. In case sex-biases in philopatry are large, selection favours near uniparental care by the most philopatric sex. Panel B: in populations with multiple breeding pairs per patch, the effect of philopatry on sex-differences in care is less pronounced. Nonetheless, even here we find that the most philopatric sex always provides the majority of care to young. Parameters: *k*_f_ = *k*_m_ = 0.1.

### 2.2 Result 2: the most common sex does not always provide more care

Next, we consider the effect of mating skew, where the number of female breeders in a patch differs from that of the male breeders, *n*_f_ ≠ *n*_m_. Note that the left-hand side of eq. (4) is multiplied by *n*_f_*/n*_m_, which shows that mating skew directly affects the equilibrium balance of sex-specific care. When males are the more common sex, for example, the left hand side in eq.(4) is reduced relative to that in eq. (3), so that an increase in male parental effort has a smaller effective mortality cost relative to an increase in female effort. Selection thus favours greater caring effort by males when they are the more common sex (and less caring effort when they are the rarer sex). That the more common sex cares more is a well-known result from studies modeling how the adult sex ratio (ASR) affects sex differences in parental care in well-mixed populations [8, 29].

However, mating skew also affects sex differences in parental effort indirectly, by modulating the relatedness coefficients (3, 4). When males are the more common sex, individual males have a lower genetic share in the next generation in the local patch (as they compete more strongly for matings), thus reducing a male’s relatedness *r*_m→locally competing juv_ to any locally competing juveniles. Vice versa, the coefficient *r*_f→locally competing juv_ for females is increased, as individuals of the rare sex have a greater genetic share in the next generation. In dying, males are therefore less likely to be replaced by a related juvenile in comparison to females, so that the effective mortality cost of care is larger for the more common sex. For the same reasons, *r*_m→competing juv_ on the right-hand side of eq. (4) is also reduced when males are the common sex, reducing the fecundity benefit of increased care for the more common sex. However, *r*_m→competing juv_ is smaller than *r*_m→locally competing juv_ on the left hand side (compare eqns. [S29, S27]), any indirect effects of mating skew via relatedness cause the more common sex to provide less care relative to the rare sex.

#### Individual care: the common sex cares more

What is the overall consequence of both indirect and direct effects on the evolution of sex-differences in care? When individuals care only for their own offspring, Figure 3A-C shows that the more numerous sex always provides the majority of care to the young. In other words, the indirect effect of mating skew via relatedness does is not enough to alter the well-established result that the more common sex cares more [8, 29]. Nonetheless, the amount of care provided by each sex is still affected by sex differences in dispersal, as levels of sex-specific care increase with philopatry (contour lines in Figure 3A-C).

**Figure 3:**
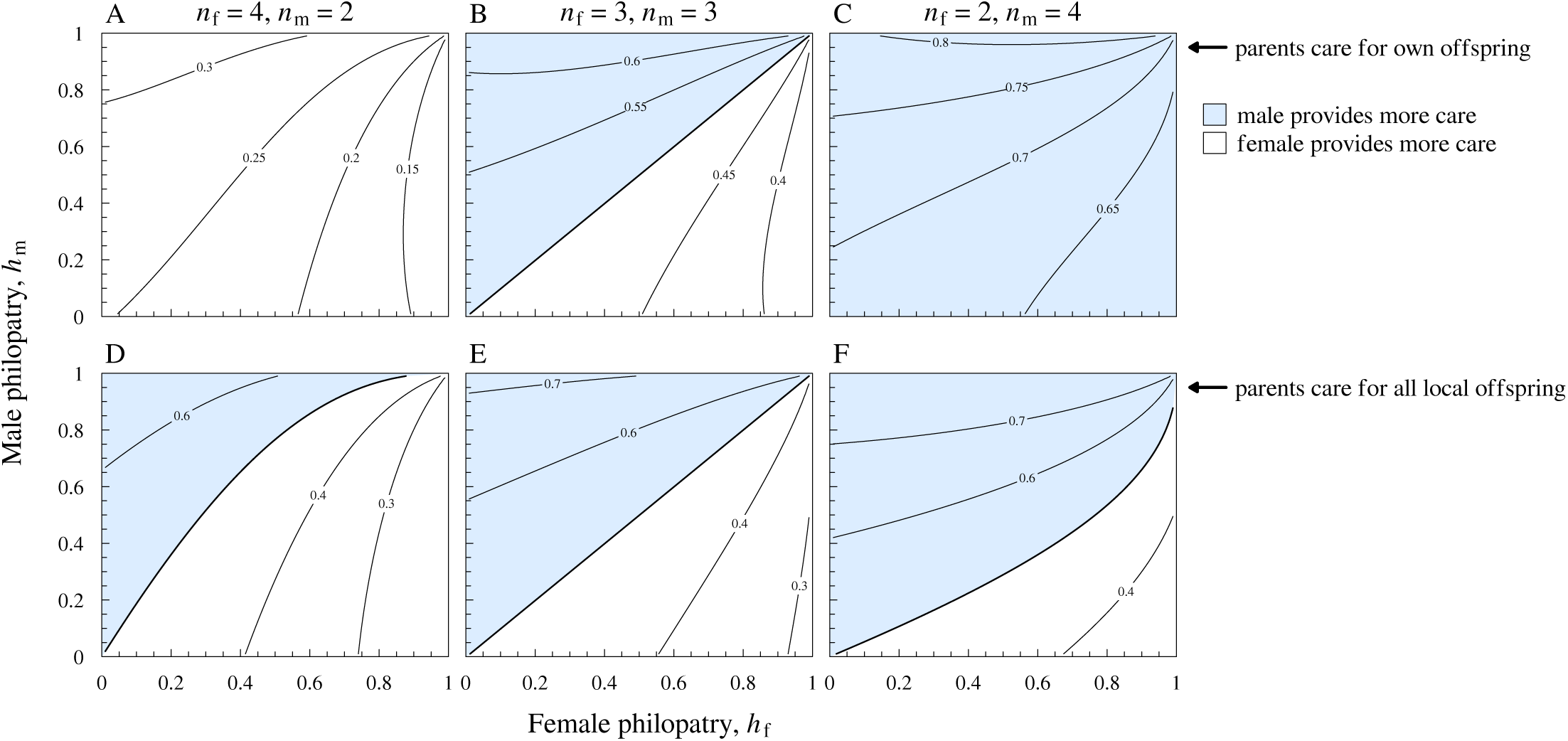
The equilibrium fraction of male care 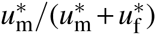 for different degrees of mating skew *n*_m_*/n*_f_. Panels A-C: Parents care only for their own offspring. When mating skews occur for parents that care only for their own offspring, the more numerous sex (i.e., the sex with the lowest reproductive value) always provides the majority of care, regardless of the level of dispersal. Nonetheless, amount of care by a particular sex always increases with its level of natal philopatry. Panels D,F: in case parents provide care for all local offspring, the parameter space in which the more numerous sex provides most of the care is substantially decreased. Consequently, the level of philopatry becomes a more important determinant of the degree of sex-specific care, so that often the philopatric sex provides the majority of care. Parameters: *k*_m_ = *k*_f_ = 0.1.

#### Communal care: the philopatric sex cares more

Things are different, however, when we consider a scenario of communal care, in which individuals provide care to all locally born offspring. Figures 3D, F show that effect of mating skew (causing the more common sex to provide more care) is now markedly reduced relative to that of sex-biased dispersal. Only when dispersal patterns are relatively similar across both sexes (i.e., close to the line *h*_m_ = *h*_f_), do we find that mating skew determines which sex provides most of the care. Otherwise, we retrieve the interesting result that the more philopatric sex provides most of the care.

Why does communal care change our conclusions? When parents care for all locally born offspring, coefficients *r*_f→own juv_ and *r*_m→own juv_ in equations (3,4) for the case of individual care are now replaced by the coefficients *r*_f→locally born juv_ and *r*_m→locally born juv_. Hence, whether focal gene copies benefit from any changes in the focal’s care now depends on mating skew: when males are the more common sex (*n*_m_ > *n*_f_), individual males have a smaller expected genetic share in the next generation relative to females. As a consequence, a slight increase in a mutant male’s parental effort is less likely to benefit copies of that mutant gene present in offspring, because *r*_m→locally born juv_ becomes smaller relative to *r*_f→locally born juv_ as *n*_m_ increases. Going back to eqns. (3,4), communal care substantially reduces the fecundity benefit of effort by the more common sex (right-hand side), thus offsetting the previously found reduction in the effective mortality cost of increased care by the more common sex (left-hand side). All in all, the effect of mating skew is therefore largely cancelled out in taxa with communal care, causing the effects of sex-biased dispersal to prevail (Figures 3D, F).

### 2.3 Result 3: sex-biased dispersal can mask sex-specific costs of care

Well-established predictions on sex-biases in care often focus on differences in costs and constraints between the sexes [17, 57, 58], which raises the question how sensitive our conclusions are to asymmetries in costs between the sexes. Figure 4 varies the mortality cost of male care relative to female care, (1 − *k*_m_)/(1 − *k*_f_). Unsurprisingly, when male philopatry is low the proportion of male care rapidly decreases with increasing mortality costs of male care. However, for taxa with high levels of male philopatry, we would predict that males still provide the majority of care in the face of considerable costs. Consequently, sex-biased dispersal can mask the effect that sex differences in cost have on the relative amount of care that males and females provide to their young.

**Figure 4:**
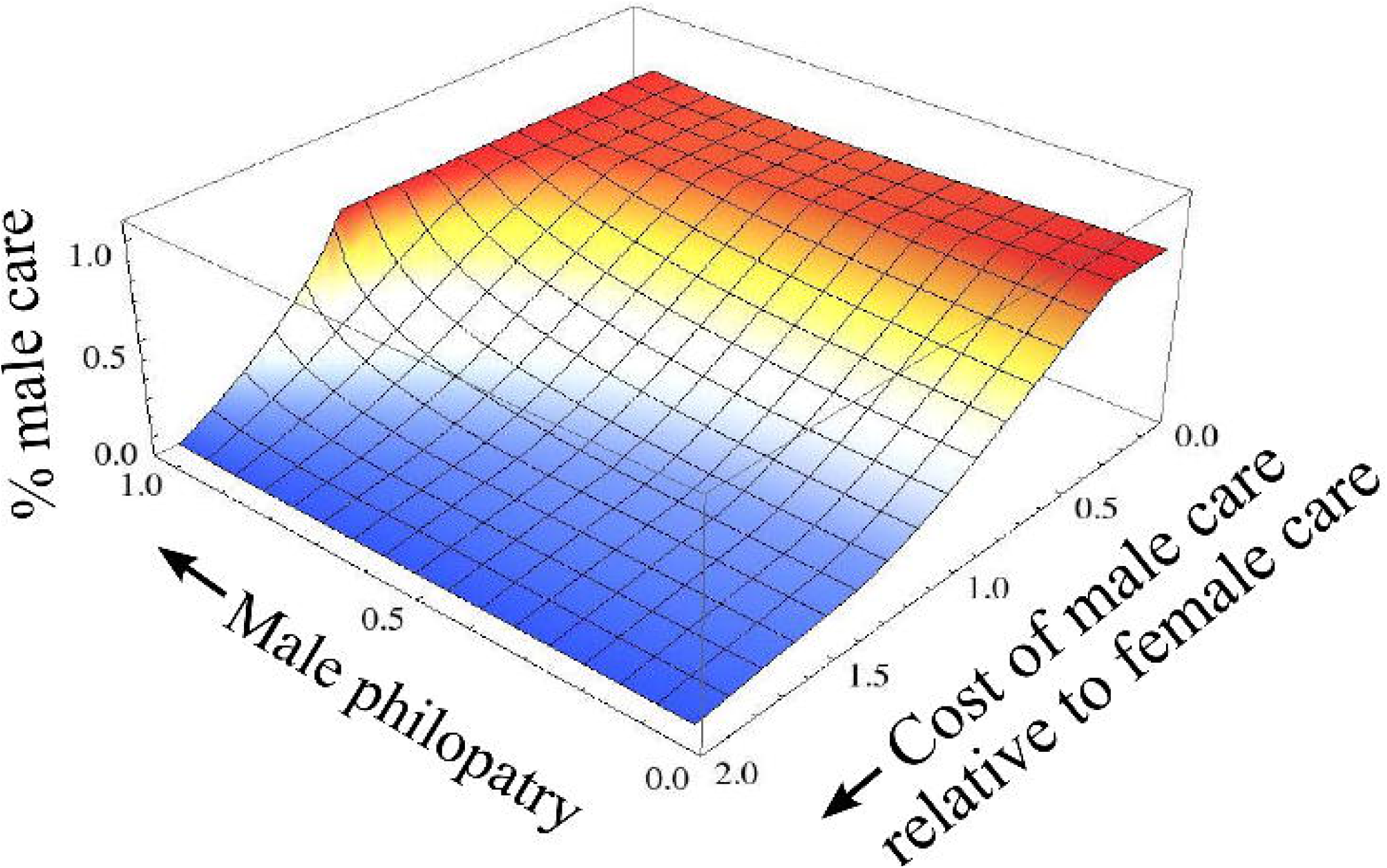
The philopatric sex (here males) typically provides the majority of care, even when it has to pay a far higher mortality cost than the dispersing sex. The equilibrium fraction of male care 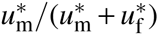 when varying the cost of male care relative to that of female care, (1 − *k*_m_)/(1 − *k*_f_) (*x*-axis) and by varying the level of male philopatry (*y*-axis). With low levels of philopatry, the proportion of male care rapidly declines from a pattern where males provide the majority of care to cases where females provide most of the care. However, when male dispersal is limited, males provide the majority of care to the brood, despite high sex-specific costs of male care. Parameters: *n*_m_ = *n*_f_ = 2, *d*_f_ = 0.1.

## 3 Discussion

Our analysis reveals that sex differences in dispersal can have a substantial effect on the resolution of sexual conflict over parental care: when females are the dispersing sex, our model predicts that males should typically invest more in caring for young. Vice versa, when males are the dispersing sex, females should invest more in care. These predictions contrast strikingly with those of classical models assuming well-mixed populations (e.g., [17]); the impact of sexbiased dispersal may even be sufficient to destabilise biparental care and lead to uniparental care by the philopatric sex (see Figure 1). Although the effect of sex-biased dispersal can be outweighed by that of mating skew in certain breeding systems (such that the more common sex provides most of the care, even if it is more prone to dispersal; Figure 3), all other things being equal, parental investment by a given sex always increases with the relative level of philopatry of that sex.

Why does sex-biased dispersal have such marked effects? Previous models of the evolution of reproductive effort in asexual populations have shown that greater effort is selectively favoured, at a cost to adult survival, when dispersal is more limited[59–61]. Our model reveals that the same holds true in populations with two sexes: where males are the philopatric sex, for example, they value the production of offspring more relative to their own survival, thus selectively favouring higher levels of male care. By contrast, the dispersing sex, in this case females, value their own survival more relative to the production of offspring, selectively favouring reduced levels of female care. These effects arise because, in dying, an adult of the philopatric sex is likely to free up a breeding vacancy that may be occupied by a local, related offspring; this indirect, kin-selected benefit partially offsets the cost of death. By contrast, when an adult of the dispersing sex dies, the vacancy created is more likely to be filled by an unrelated, immigrant offspring. Consequently, as sex-biased dispersal changes the trade-off between adult survival and offspring production to diverge between the sexes, sex-biases in parental care are likely to arise.

Our model also shows that mating skew can substantially influence sex-specific patterns of parental care, with the more numerous sex (all other things being equal) providing more of the care. This finding corroborates previous studies on the relationship between the ASR and sex roles in well-mixed populations [8, 28, 29]. As one sex becomes more numerous, investment in survival becomes less advantageous due to increased competition over future breeding opportunities, favouring increased investment in current reproductive output. Consequently, we expect the more numerous sex to provide most of the care regardless of the degree of sex-biased dispersal (e.g., Figure 3A-C). However, the same does not hold in populations with communal care, in which local males and females care for all locally born offspring. In the case of communal care, parental investment by individuals of the more numerous sex yields fewer benefits because care is distributed among offspring other than one’s own, favouring lower levels of investment.

Our prediction that the philopatric sex provides more care could be tested using available data on parental care and sex-biased dispersal in well-studied taxonomic groups. One may, for instance, contrast dispersal and parental care in birds and mammals: in mammals, males are typically the dispersing sex [37, 39, 62] and indeed we find that the more philopatric sex (females) provide most of the care: in 90% of all mammalian taxa female care predominates, while male-only care is completely absent [63]. In birds, by contrast, females are typically the sex more prone to dispersal [37, 38, 64], and male contribution to parental care is much more common in birds than in mammals, with the vast majority of all bird species (90%) exhibiting biparental care [63]. Obviously, there are other differences between birds and mammals that might account for these broad differences, and a more detailed comparative analysis would be needed to test our predictions properly. Nevertheless, our model clearly suggests that spatial structure and sex-differences in dispersal have been largely overlooked in existing meta-analyses (e.g., [65–67]) that aim to understand the social and ecological factors influencing sex role evolution.

Previous models have focused on behavioural interactions at the level of the family and on sex-specific costs of care as the main drivers of asymmetries in parental care between the males and females [17–23, 68, 69]. To our knowledge, there are currently no theoretical predictions that take account of ecological processes at population-level. Since our results show that dispersal and spatial population structure affect the outcome of conflict between parents over care, it seems likely that other ecological factors may also have a hitherto unforeseen impact on the relative level of investment by mothers and fathers. Factors such as spatiotemporal variation in resources, predation and disease are likely to be important factors [70], but the impact of such variation on parental care tactics remains to be explored.

Possible extensions to our model include analysing the coevolution of care and dispersal strategies. We have treated sex-specific dispersal rates as fixed parameters, and yet the evolution of sex-biased dispersal itself may well be affected by male and female parental effort. So far, studies of the evolution of sex-differences in dispersal have focused on inbreeding avoidance and sex differences in kin competition as drivers of sex-biases in dispersal [71–74],. overlooking the impact of sex-differences in care. In the absence of any sex-specific fitness consequences of parental effort, investment in care should affect competition among kin of both sexes equally and thus is unlikely to lead to sex-differences in dispersal. However, when parental care is more beneficial for offspring of one sex versus the other (e.g., [75]), parental effort is likely to affect sex-biases in dispersal.

Another key assumption of our model is that parental strategies specify investment in care as a ‘sealed bid’ by both parents, as in the classical model by Houston & Davies [17]. In other words, parents are unable to respond to each other’s efforts on a behavioural time scale. By contrast, more recent models of negotiation over care have begun to explore the evolution of sex-differences in the degree of responsiveness to the amount of effort invested by a partner [18, 19, 21, 22]. When responding to each other’s efforts is costly, our study suggests that sex-biases in dispersal are likely to affect the degree of sex-specific responsiveness, simply because the effective cost of being more responsive will be lower for the philopatric sex. In general, sex differences in phenotypic plasticity (of which parental responsiveness is an example) are likely to be affected by the degree of philopatry.

To summarise, more studies that place models of family conflict in their population-level, ecological context are likely to enhance our understanding of the great variation in family life histories that we observe in nature.

## Acknowledgements

We would like to thank the other members of the Transgen group, Tom Ezard, Stuart Townley and Jonathan Wells for discussion. The Dutch Academy of Arts and Sciences (KNAW) and the Lorentz Centre at the University of Leiden, the Netherlands, funded a week-long workshop on nongenetic effects that contributed to this paper. The authors acknowledge the use of the UCL Legion High Performance Computing Facility (Legion@UCL), and associated support services, in the completion of this work. This study was funded by an EPSRC sandpit grant on transgenerational effects, grant number EP/H031928/1 awarded to RAJ and an EPSRC-funded 2020 Science fellowship awarded to BK (grant number EP/I017909/1).

